# Neuritin1 *Cis*-Regulatory Elements Enable Gene Expression Preferentially in Retinal Ganglion Cells

**DOI:** 10.64898/2026.04.01.715961

**Authors:** Venu Talla, Rajeshwari Koilkonda, Chaimaa Kinane, Moxa Panchal, Tamdan Khuu, Kevin K. Park

## Abstract

**Purpose:** Retinal ganglion cells (RGCs) are essential for visual signal transmission, yet they are vulnerable to injury and degeneration. Gene modulation in RGCs using adeno-associated virus (AAV) offers a promising avenue for neuroprotection and regeneration, but promoters lack sufficient RGC specificity, limiting precision needed for preclinical studies. This study aims to identify novel promoter-enhancer combinations (PECs) to achieve gene expression preferentially in RGCs.

**Methods:** We evaluated existing transcriptomic data to identify Neuritin 1(Nrn1) as a gene with highly restricted RGC expression in the retina. Synthetic PECs derived from human and mouse Nrn1 loci were incorporated into AAV2 vectors driving expression of a nuclear-targeted reporter GreenLantern. AAVs were delivered via intravitreal injection into C57BL6/J mice, and transduction efficiency and RGC specificity were evaluated in both young and aged retinas and those subjected to intraorbital optic nerve crush (ONC), using immunohistochemistry and quantitative analysis of RBPMS+ cells.

**Results:** We found that AAV2 with a human Nrn1 PEC drives gene expression in RGCs. Quantitative analysis revealed that over 83% of transduced cells were RBPMS-positive, indicating robust RGC selectivity and significantly outperforming ubiquitous promoters. Notably, the Nrn1 PEC retained strong and selective transgene expression in RGCs in aged mice and following ONC, demonstrating its resilience under aged and injury conditions.

**Conclusion:** The Nrn1 PEC enables efficient and injury-resilient gene expression in RGCs, addressing a key limitation in cell-specific targeting. This AAV-incorporated PEC offers a robust platform for evaluating neuroprotective interventions and accelerates translational development of gene therapies for glaucoma and other optic neuropathies.

## INTRODUCTION

Retinal ganglion cells (RGCs) are the sole output neurons of the retina, transmitting visual information from photoreceptors to the brain through the optic nerve. They are essential for conscious vision, yet their loss is irreversible, as RGCs do not regenerate in humans. Degeneration of RGCs underlies multiple blinding conditions, including glaucoma, optic neuritis, ischemic optic neuropathies, and traumatic injury. Despite decades of research, no therapies reliably prevent RGC loss or restore function^1–9^, representing a profound unmet need in ophthalmology and a critical barrier to preserving vision.

Gene therapy holds significant promise for protecting RGCs, promoting axonal repair, and mitigating disease-specific insults. However, many therapeutic candidates act cell-autonomously within retina and exert broad biological effects that can disrupt retinal circuitry or cause harm when expressed in non-RGC cell types. Therefore, achieving RGC-specific gene manipulation is critical for both safety and efficacy. This need is underscored by a key technical limitation: most promoters used in adeno-associated virus (AAV) vectors lack sufficient specificity for RGCs. Robust, RGC-selective targeting is essential to advance preclinical studies and enable precise, effective gene therapies for clinical translation.

Existing promoters designed for RGCs enhance RGC selectivity compared to ubiquitous promoters.^10–13^ However, many remain suboptimal, particularly at the high viral titers required for efficient transduction. At low viral titers, these promoters drive expression predominantly in RGCs; however, at higher titers, high percentage of the transduced cells are non-RGCs.^12^ This presents a fundamental trade-off: achieving broad RGC transduction comes at the cost of reduced specificity, whereas maintaining selectivity limits transduction efficiency. This limitation constrains both preclinical studies and future translational applications, underscoring a critical technical and conceptual gap in the field.

To address this need, we performed analysis on the existing single-cell transcriptomics data of mouse retinas, identifying Neuritin1 (Nrn1) as a gene with highly restricted RGC expression, highlighting its potential as a cell type–specific regulatory element. To determine whether Nrn1 cis-regulatory elements can be leveraged to drive transgene expression in RGCs, we screened multiple promoter and enhancer combinations (PECs) from both human and mouse Nrn1 genes. Through this work, we demonstrate that AAV2 incorporating a human Nrn1 PEC achieves preferential gene expression in RGCs following intravitreal injection. Moreover, Nrn1 PEC confers strong RGC preference and transgene expression even in aged animals or following optic nerve crush (ONC). The combined properties of injury resilience, age stability, and cell-type specificity position the Nrn1 PEC as a superior regulatory element for gene modulation strategies targeting RGCs.

## METHODS

### Construction of Nrn1-monomeric Green Lantern (mGL) AAV Vectors

Three promoter enhancer combination (PEC) sequences were synthesized as gene fragments by Integrated DNA Technologies (IDT, Coralville, IA, USA). Each fragment contained 25-nt homology arms corresponding to regions flanking the XbaI and HpaI sites in the pAAV-CMV-H2b-mGL vector. The vector was digested with XbaI and HpaI, and the 4,286-bp backbone was gel-purified. The gel purified vector back bone was combined with PEC gene fragment and pAAV-Nrn1PEC constructs were assembled using the NEB HiFi DNA Assembly Master Mix (New England Biolabs, Ipswich, MA; Cat. #E2621). Assembled plasmids were introduced into NEB Stable competent cells (New England Biolabs, Ipswich, MA), and positive clones, including ITR integrity, were verified by restriction analysis and whole-plasmid sequencing.

### AAV Production and Purification

AAVs were produced using a standard triple-transfection approach in VPC 2.0 suspension cells (Thermo Fisher Scientific). Briefly, cells were seeded one day prior to transfection. On the following day, cultures were adjusted to 3 × 10^6^ cells/mL and transfected with pHelper (Aldevron), pRepCap2, and pAAV-Nrn1-PEC plasmids at a 1:1:1 ratio. Polyethylenimine (PEI) was used as the transfection reagent at a 2:1 PEI:DNA mass ratio. Three days after transfection, cells were harvested by centrifugation and lysed using three cycles of freeze–thaw. The crude lysate was incubated with Benzonase (50 U/mL) at 37 °C for 30 min with gentle shaking to degrade contaminating host DNA and RNA. Lysates were then clarified by centrifugation at 4,000 × g for 20 min at 4 °C, followed by filtration through a 0.2 μm membrane. The clarified material was loaded onto an AAVX Affinity POROS GoPure FPLC column for capture of AAV particles. To further purify, the eluted material underwent iodixanol gradient ultracentrifugation. Final viral titers were quantified using the Applied Biosystems Absolute Q™ Viral Titer dPCR Assay (Thermo Fisher Scientific, Cat. No. A53736) on the Applied Biosystems QuantStudio Absolute Q™ Digital PCR System (Thermo Fisher Scientific). The final titers of AAVs ranged from 1-3 x 10^13^ genomic copies (gc)/ml.

### Experimental Animals and Intravitreal Injection

All animal procedures were performed in compliance with the protocols approved by the Institutional Animal Care and Use Committee at University of Texas Southwestern Medical Center and in accordance with the ARVO Statement for the Use of Animals in Ophthalmic and Vision Research. For the intraocular injection of recombinant AAV, 6-8wk old (young) or 8-month-old (aged) male C57BL/6J mice (Jackson Lab, Stock# 000664) were sedated by inhalation with 1.5% to 2% isoflurane and a 33-gauge needle attached to a Hamilton syringe was inserted through the pars plana. Left eyes were injected with about 1.5 ul of AAV2-Nrn1-H2B-mGL or AAV2-CMV-H2B-mGL to give a final dose of approximately 1.5 x 10^10^ gc/injection. Contralateral right eyes remained as the un-injected controls. Following the injections, animals received subcutaneous injection of Buprenorphine as an analgesia. Numbers of animals used were two to three per treatment group. The numbers of animals used are indicated in each figure legend.

### Optic nerve crush (ONC)

ONC was performed three weeks following the AAV injections. For this, 6-8wk old male C57BL/6J mice were sedated by inhalation with 1.5-2% isoflurane, optic nerve was exposed intraorbitally without damaging underlying ophthalmic artery and crushed using a jeweler’s forceps (#5 Dumont, Fine Science Tools) for 15 seconds approximately 1 mm behind the eyeball. Following the ONC, animals were administered subcutaneous injection of Buprenorphine as an analgesia.

### Retina Preparation for Flat mount and Cryosection

Three weeks post AAV injection, animals were transcardially perfused with cold phosphate buffered saline (PBS, pH 7.4) until blood was removed from the system, followed by the administration of ∼25 ml of 4% paraformaldehyde (PFA, pH 7.4). Eyes were enucleated and post-fixed for 2-3hrs in 4% PFA at room temperature. Tissue was then washed with PBS (1X) and transferred to 30% sucrose in PBS for cryoprotection, remaining at 4°C overnight until the tissue sank. For longitudinal sections eyes and retinas were embedded in optimal cutting temperature, frozen on dry ice, and stored at −80°C until use. Retinas were sectioned at 10 µm using cryostat (Leica CM3035) and mounted on SuperfrostPlus slides. Slides were air-dried for about 30-40min and stored at −20°C until immunostaining.

### Immunohistochemistry

For immunohistochemistry, retinas were dissected out for whole-mount, transferred to the 24 well plate and were blocked in 2% Triton X-100 and 10% goat serum in PBS for 1hr. Retina whole mounts were incubated with a mixture of guinea pig anti-RNA Binding Protein, MRNA Processing Factor (RBPMS) antibody (1:2000, ProSci, custom generated) and chicken anti-GFP (i.e., mGL) antibodies (1:200, ThermoFisher Scientific, Cat#A10262) overnight at 4°C. For the retinal sections, the same staining protocol was used on the slides. Retinas were washed with PBS and incubated in secondary antibody solution containing goat-anti guinea pig DyLight 800 (1:500, ThermoFisher Scientific, Cat#SA5-10100) and Goat anti-chicken Alexa Fluor 488 (1:500, Jackson ImmunoResearch Laboratories, Cat#103-545-155) at room temperature for 1hr. Following three washes with PBS, retina whole mounts were mounted using Vectashield antifade mounting media (Vector Laboratories). Fluorescence images were acquired using a Keyence BZX-800 microscope. For RGC counts on retina whole mounts, ten images were sampled around the central, mid periphery and peripheral regions of each retina whole mount (0.1 µm^2^ area/image). For longitudinal sections Z-stack images focusing RGC somas followed by processing to generate a composite image was performed. As RGC density may vary across the retina, five Z-stack images were taken across the entire retina section. Multiple sections and 2-3 slides were used to image each sample. Of note, although mGL is engineered to be brighter and more stable than GFP, we used anti-GFP antibody staining to ensure sensitive detection of mGL signal. This approach minimizes the risk of underestimating transgene expression due to variability in expression level, tissue depth, or fixation-related signal loss, and allows for more consistent and unbiased quantification across samples.

### Quantitative Analysis

ImageJ software was used for quantitative analysis of mGL positive cells that are RBPMS positive or RBPMS negative in the retina flat mounts or longitudinal sections. Statistical analyses were performed using GraphPad Prism 10. Data are presented as mean±SD. One-way ANOVA followed by Tukey’s post hoc test or Welch’s t test was used and p<0.05 was considered statistically significant.

## RESULTS

### Nrn1 promoters and enhancers as candidate cis-regulatory elements (CREs) to drive gene expression in RGCs

Cell type–specific promoters or enhancers can be incorporated into adeno-associated virus (AAV) vectors to confine transgene expression to target cells. To identify CREs capable of driving gene expression specifically in RGCs, we analyzed single-cell RNA-seq data from the healthy adult mouse retina.^14^ This analysis revealed that *Nrn1* is expressed almost exclusively in RGCs whereas other top RGC markers gamma synuclein (*Sncg*), *Thy1*, neurofilament light (*Nefl*), stathmin-2 (*Stmn2*), *Rbpms*, and ubiquitin C-terminal hydrolase L1 (*Uchl1*) are still expressed at considerable levels in one or more retinal cell types (Supplementary Fig. 1A). *Pou4f1* expression is specific to RGCs but its expression likely diminishes after injury.^15^ We observed from the RGC scRNAseq that *Nrn1* is expressed across all RGC subtypes (Supplementary Fig. 1B), and that its expression does not decrease drastically soon following optic nerve crush (Supplementary Fig. 1C).^15^

We searched for the presence of promoter and enhancer elements in proximity to the Nrn1 gene using the data from UCSC Genome Browser and ENCODE (Encyclopedia of DNA Elements).^16^ Candidate regions were prioritized based on ReMap transcription factor binding density and FANTOM5 CAGE (Cap Analysis of Gene Expression) peaks indicative of transcription initiation^17^ (Supplementary Figs. 2A and 2B). ReMap density plots visualize transcription factor binding across cell types using aggregated ChIP-seq data. They highlight regulatory hotspots—regions with high transcription factor occupancy—suggesting potential enhancers, promoters, or other functional elements. Three distinct promoter–enhancer combinations (PECs) were selected from the human and mouse genomes for synthetic sequence testing (Supplementary Figs. 2A and 2B). Each PEC was cloned into an AAV vector containing a nuclear-targeted mGL reporter to assess cell type-specific regulatory activity (Figs. 1A-C).

**Figure 1.**
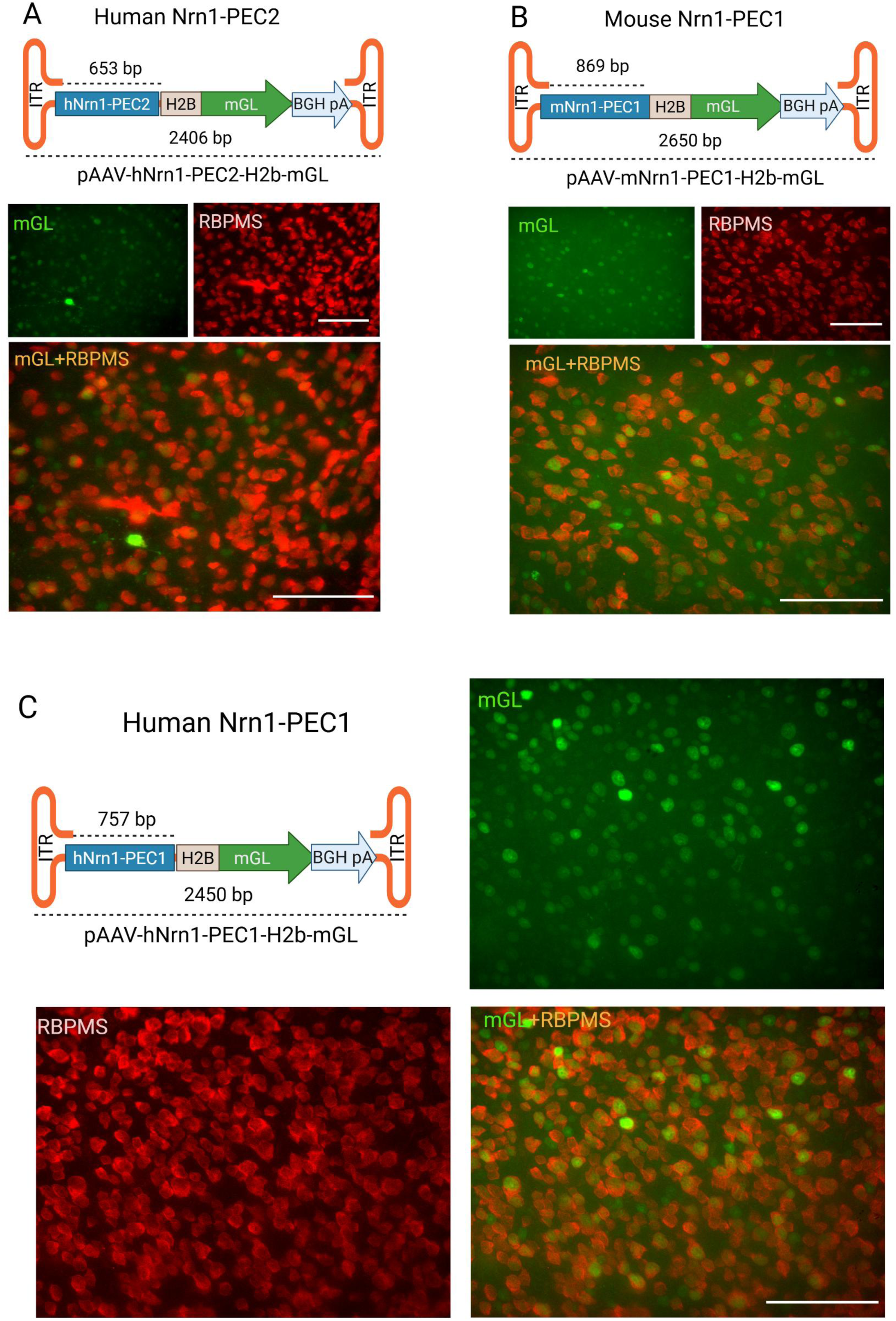
(**A-C**) Schematic diagrams and representative retinal fluorescence images showing nuclear mGreenLantern (mGL, green) expression driven by hNrn1-PEC2 (A), mNrn1-PEC1 (B), and hNrn1-PEC1 (C) promoters in AAV constructs. The lengths of the PEC sequences and the total size of each recombinant AAV genome are indicated in the schematics (e.g., for hNrn1-PEC1, the length of PEC is 757bp and the entire length between the two ITRs is 2,450bp). For the flat mounted retinal images, red signal marks RBPMS immunoreactivity. Images were obtained from retinas subjected to three weeks post AAV injection. N=3 per group.

**Figure 2.**
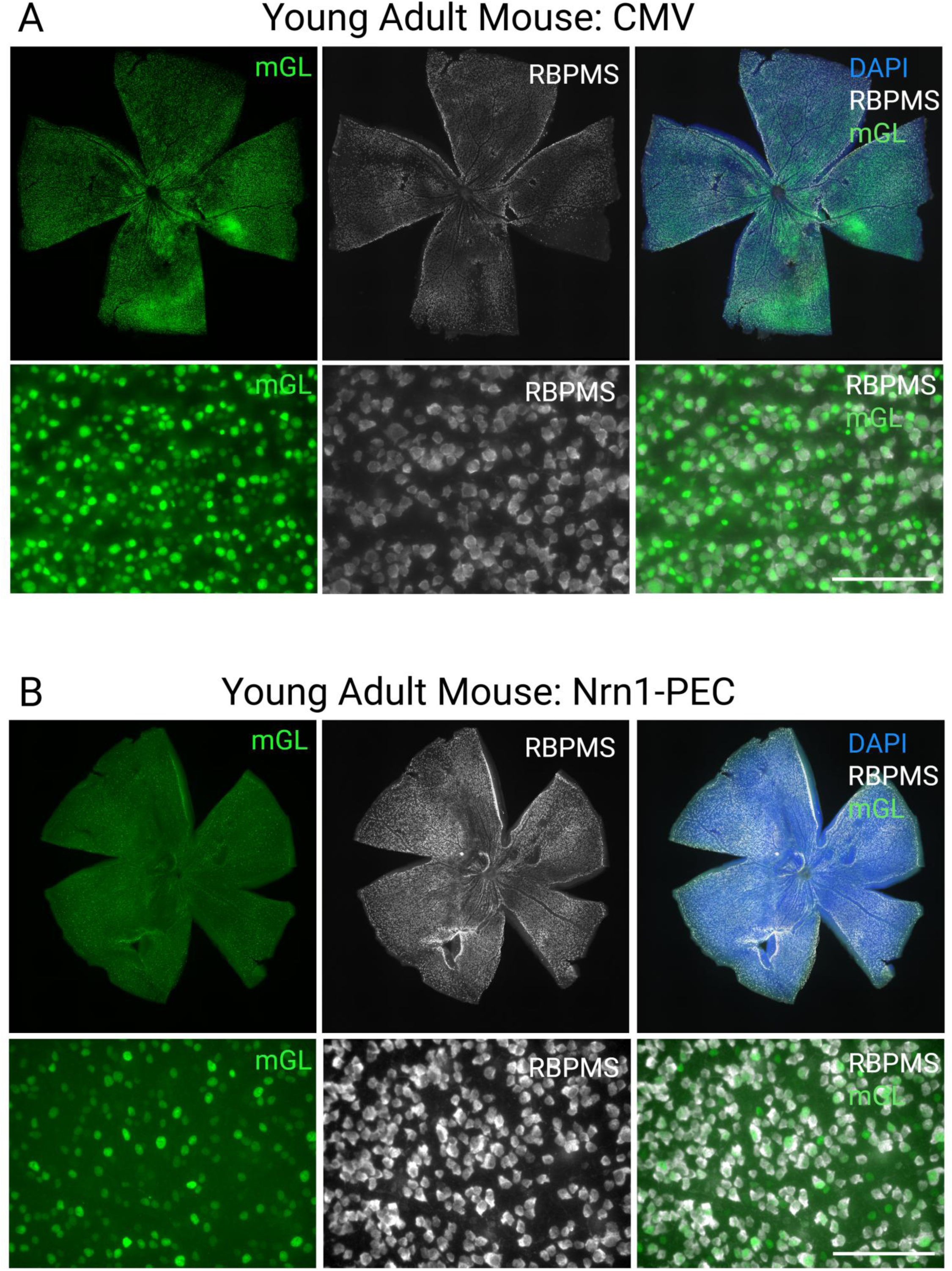
Nrn1 PEC drives strong reporter expression in RGCs across the retina following intravitreal injection. (**A**) Representative retinal wholemounts and high-magnification images showing mGL fluorescence from an AAV2-CMV animal. (**B**) Representative retinal wholemounts and high-magnification images showing mGL fluorescence from a Nrn1 PEC1 animal. mGL expression colocalizes with the RGC marker RBPMS. Scale bars, 100µm.

To compare transcriptional activity among the human and mouse Nrn1 PEC variants, young adult mice (6-8 weeks old; n = 3 per group) received intravitreal injections of AAVs expressing H2B-mGL under the control of either human Nrn1 (hNrn1)-PEC1, hNrn1-PEC2, or mouse Nrn1 (mNrn1)-PEC1 regulatory elements. Three weeks after administration, retinal whole mounts showed nuclear mGL expression in RBPMS⁺ RGCs across all groups, with distinct differences in signal intensity and uniformity. The hNrn1-PEC1 construct (Fig. 1C) produced strong nuclear labeling and yielded the highest density of mGL labeled RGCs. In contrast, the shorter hNrn1-PEC2 promoter (Fig. 1A) drove lower overall fluorescence and fewer strongly expressing cells, indicative of weaker promoter activity. The mNrn1-PEC1 promoter (Fig. 1B) produced expression levels higher than hNrn1-PEC2 but remained weaker and less uniform than hNrn1-PEC1. Together, these findings demonstrate that promoter length and enhancer composition modulate Nrn1-driven reporter expression in the retina, with hNrn1-PEC1 (hereafter referred simply as Nrn1 PEC) exhibiting the strongest activity and selected for further characterization.

### Nrn1 PEC drives gene expression preferentially in RGCs following intravitreal injection

To assess the cell-type specificity of the Nrn1 PEC upon high-titer AAV administration, mice received two intravitreal injections of AAV2-hNrn1PEC1-H2B-mGL or AAV2-CMV-H2B-mGL spaced one day apart, and retinal wholemounts were examined three weeks after the first injection. The Nrn1 PEC drove robust nuclear mGL expression that colocalized with RBPMS-positive RGCs, with minimal expression in non-RGCs (Fig. 2B). In contrast, AAV2-CMV-H2B-mGL produced non-selective expression across both RGC and non-RGC populations (Fig. 2A). Low-magnification images showed clear reporter expression across the retina, consistent with improved distribution from two injection sites. To further characterize the identity of mGL cells, we immunostained retinal sections (Figs. 3A and 3B). Quantification demonstrated that approximately 47% of RBPMS positive RGCs in the retina exhibited mGL expression in the Nrn1 PEC group (Figs. 3C and 3E). On the other hand, CMV group showed that 92% of RGCs express mGL even though many non-RGCs also express mGL (Figs. 3C and 3E).

**Figure 3.**
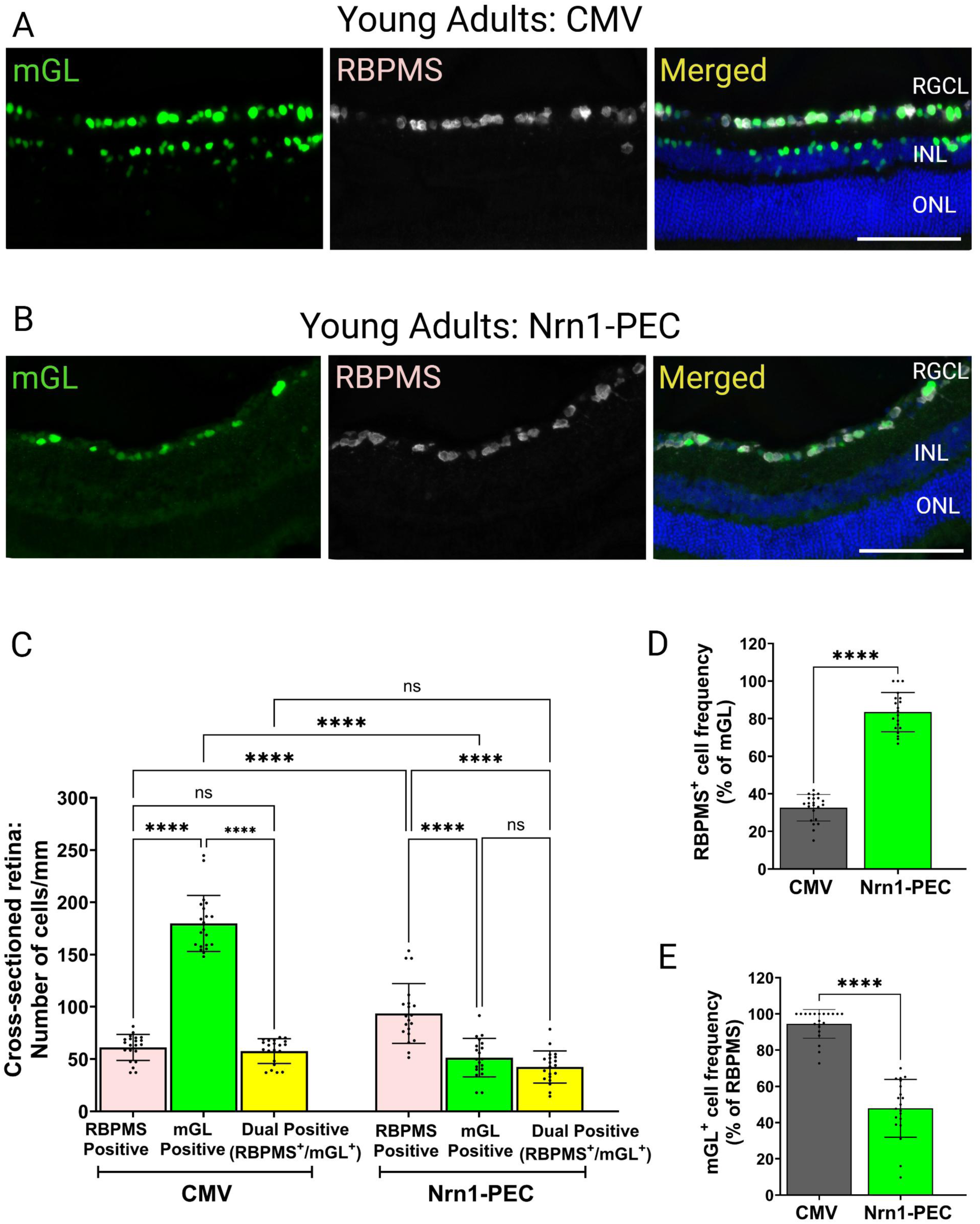
Nrn1 PEC drives gene expression preferentially in RGCs following intravitreal injection. (**A**) Retinal section from an AAV2-CMV animal. CMV drives strong mGL expression in both the ganglion cell and the inner nuclear layers. (**B**) Retinal sections from Nrn1 PEC1 animal showing mGL expression restricted in the ganglion cell layer. INL, inner nuclear layer; ONL, outer nuclear layer; RGCL, retinal ganglion cell layer. Scale bar, 100µm. N=2 per Nrn1 PEC group; N=2 per AAV-CMV group. (**C**) Quantification of RBPMS⁺, mGL^+^ and RBPMS⁺mGL⁺ cells per mm demonstrates the strength of these promoters. (**D**) RBPMS⁺ frequency (% of mGL⁺) as a measure of RGC specificity. Approximately 32% and 83% of mGL^+^ cells expressed RBPMS in the AAV-CMV and AAV-Nrn1-PEC-injected retinas, respectively. (**E**) mGL+ frequency (% of RBPMS) as a measure of transduction efficiency in RGCs. Approximately 92% and 47% of RBPMS^+^ cells expressed mGL in the CMV and Nrn1-PEC injected retinas, respectively. N= 3 per group. The scatter plots represent retinal section images. Data are mean ± SD; One-way ANOVA followed by Tukey’s multiple comparison test for graph C and Welch’s t test for graph D and E.****p < 0.0001, ns = not significant.

Importantly, our findings demonstrate that the human Nrn1 PEC confers markedly enhanced selectivity for RGCs compared to the widely used CMV promoter. In the retinal sections of CMV animals, we detected approximately 175 mGL⁺ cells/mm across all retinal layers, of which 55 were RBPMS positive and ∼120 were RBPMS negative, indicating that ∼32% of mGL positive cells are RGCs. In contrast, in Nrn1 PEC retinas, we observed ∼50 mGL positive cells/mm, of which ∼42 cells were RBPMS positive and the remainder RBPMS negative (Fig. 3C). Thus, this analysis reveals that over 83% of mGL cells were RGCs with the Nrn1 PEC, indicating strong preference for RGCs (Fig. 3D). In these Nrn1 PEC retinas, we also observed small populations of mGL positive cells that appeared RBPMS-negative, located both in the GCL and the inner nuclear layer (INL). No mGL positive cells were observed in the outer nuclear layer or the retinal pigment epithelium (Fig. 3B). Together, these data demonstrate that while the CMV promoter drives a higher overall number of mGL positive cells than the Nrn1 PEC, this increased transduction efficiency comes at the expense of reduced RGC specificity.

### Nrn1 PEC drives gene expression preferentially in RGCs in aged retina

Retinal ganglion cell degeneration is a hallmark of numerous age-related ocular diseases, including glaucoma, optic neuritis, and ischemic optic neuropathy. As such, aged mouse models are frequently employed to study disease mechanisms and evaluate therapeutic strategies that target RGCs. To assess the performance of the human Nrn1 PEC under these clinically relevant conditions, we conducted intravitreal injections with the dose and volume as mentioned above in 8-month-old mice. Strikingly, the Nrn1 PEC retained its high degree of RGC specificity in the aged retina (Fig. 4A). Immunohistochemical analysis revealed that over 83% of transduced cells were positive for RBPMS (Fig. 4B). This level of selectivity was comparable to that observed in younger adult retinas (Supplementary Figure 3), indicating that the targeting fidelity of the Nrn1 PEC is preserved with age.

**Figure 4.**
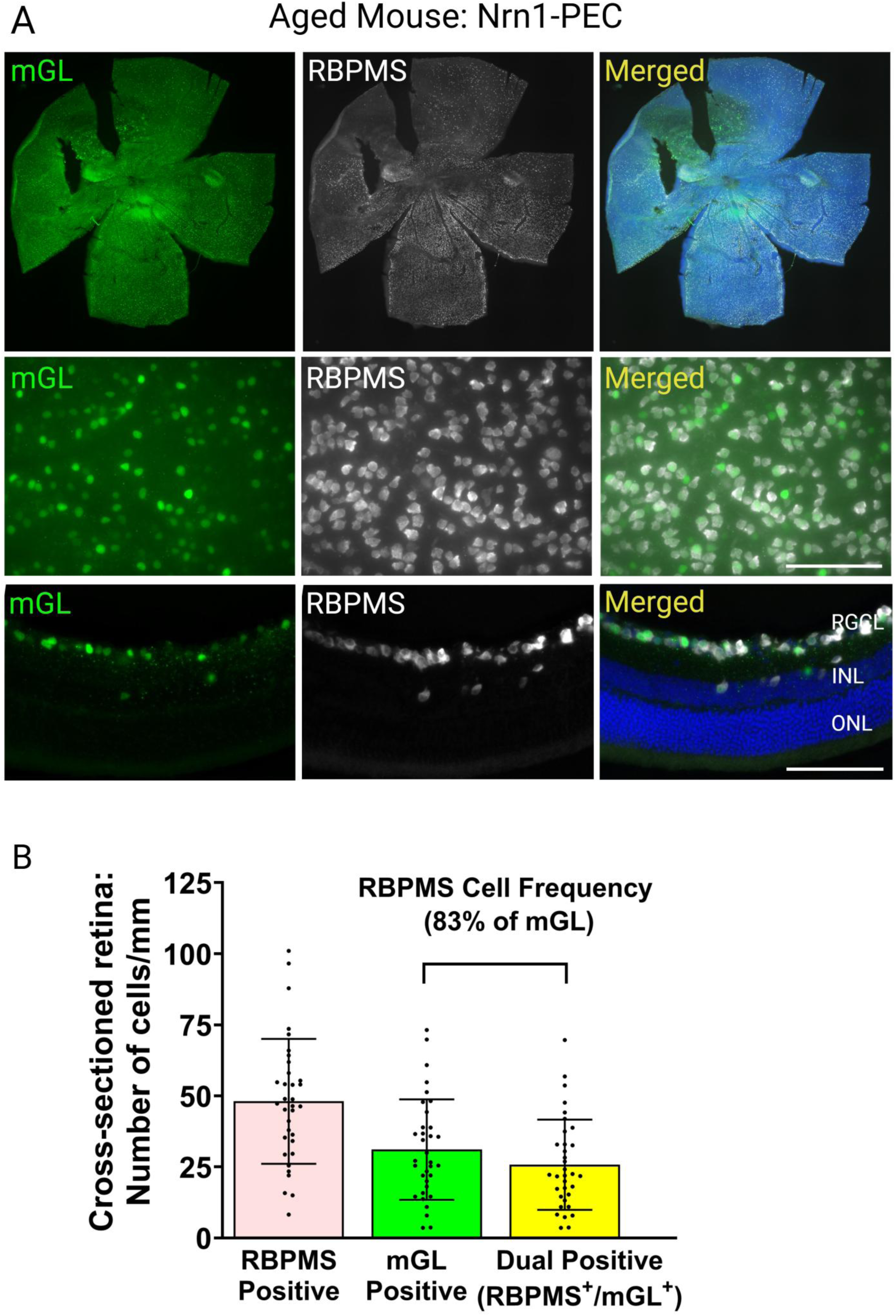
Nrn1 PEC drives gene expression preferentially in RGCs in aged mice. (**A**) Representative fluorescence images of retinal whole mounts at a low magnification (top), at a higher magnification (middle), and cross sections (bottom) from aged mice injected with AAV-Nrn1-PEC, examined three weeks post-injection. Images show mGL fluorescence (green), RBPMS (gray), and merged channels with DAPI-labeled nuclei. Scale bar equals to 100µm. (**B**) Quantification of RBPMS⁺, mGL^+^ and RBPMS⁺mGL⁺ cells per mm. Data are mean ± SD. N= 2 per group. The scatter plots represent retinal section images.

### Nrn1 PEC retains efficient and preferential gene expression in RGCs following optic nerve crush

Optic nerve crush (ONC) is a widely used model of traumatic axonal injury that mimics key pathological features of glaucoma and other optic neuropathies, including progressive RGC degeneration^18–23^. To determine whether the human Nrn1 PEC remains effective under conditions of acute neuronal stress, we performed intravitreal injections as above in young adult mice followed by ONC three weeks later. Retinas were assessed 3 days after ONC. Despite the injury-induced onset of RGC degeneration, the Nrn1 PEC maintained transduction efficiency and cell-type specificity (Fig. 5A). Immunohistochemical analysis revealed that over 78% of transduced cells were positive for RBPMS, confirming selective gene expression in the injured RGCs (Fig. 5B). Thus, the results from this and the preceding experiment demonstrate that the level of specificity in aged and ONC retinas is maintained relative to uninjured controls (Supplementary Figure 3).

**Figure 5.**
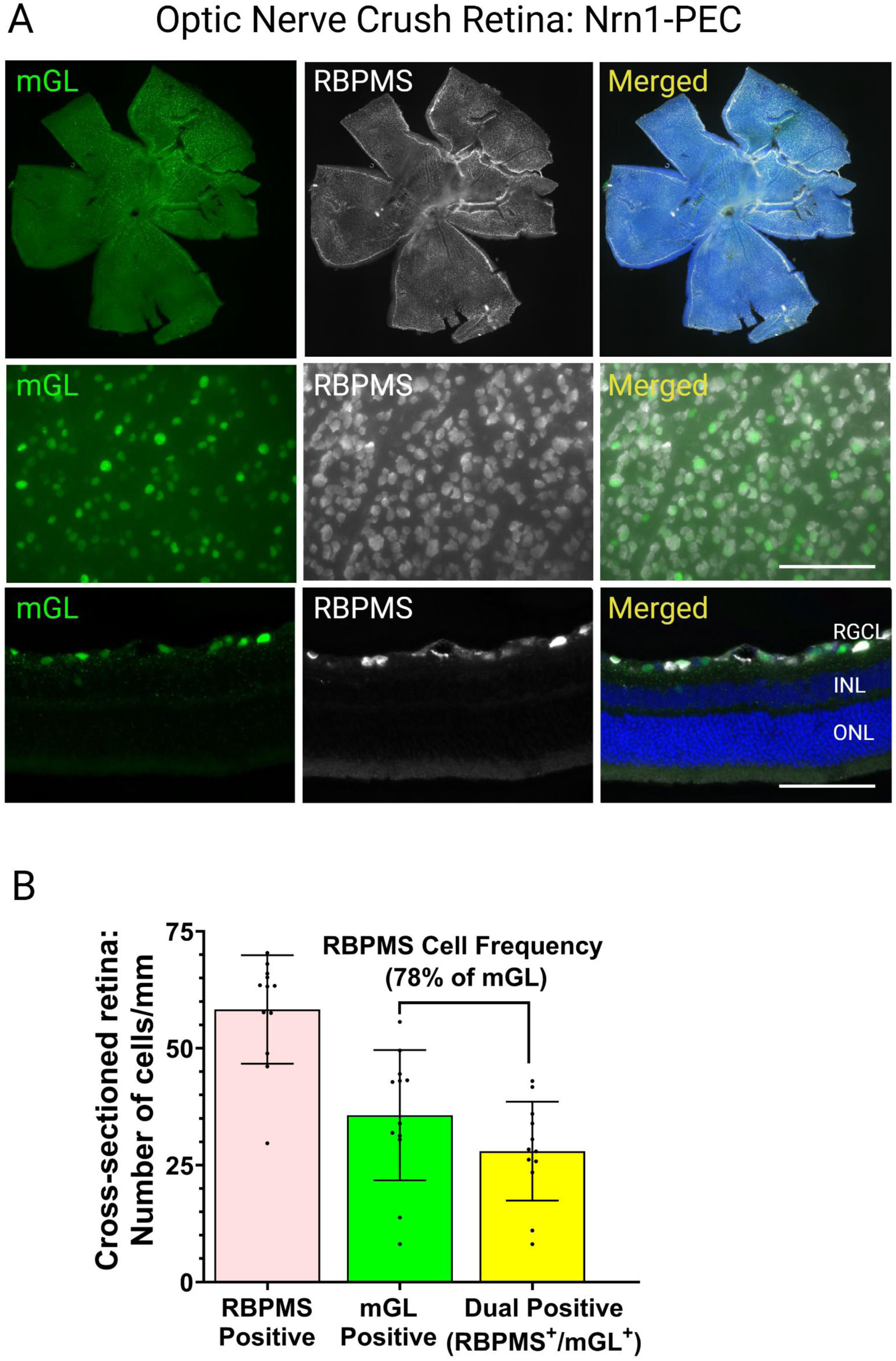
Nrn1 PEC drives gene expression preferentially in RGCs after optic nerve crush. (**A**) Representative fluorescence images of retinal whole mounts at a low magnification (top), at a higher magnification (middle), and cross sections (bottom) from ONC mice injected with AAV-Nrn1-PEC, examined 3 weeks post-injection. Images show mGL fluorescence (green), RBPMS (gray), and merged channels with DAPI-labeled nuclei. Scale bar equals to 100µm. (**B**) Quantification of RBPMS⁺, mGL^+^ and RBPMS⁺mGL⁺ cells per mm. Data are mean ± SD. N= 2 per group. The scatter plots represent retinal section images.

## DISCUSSION

Gene therapy holds significant promise for preserving RGCs or regenerating their injured axons. However, it is plausible that some neuroprotective genes act cell-autonomously in RGCs yet can be harmful if expressed in other retinal cells, causing disrupted synaptic connectivity, degeneration, or inflammation. For example, studies have shown that silencing the tumor suppressor and RGC death–promoting factor p53 in Müller glia leads to aberrant proliferation, raising the risk of tumor formation.^24^ Similarly, NFκB overexpression protects RGCs, but ectopic expression in non-RGCs induces high levels of TNFα, IL-1β, and chemokines, promoting neuroinflammation and excitotoxicity.^25,26^ These examples illustrate how off-target gene expression can cause unintended toxicity or disruption of retinal circuitry, emphasizing the need for highly selective RGC-targeted strategies to ensure both safety and therapeutic efficacy.

Many previous studies have used ubiquitous promoters, such as CAG and CMV, because of their strong ability to transduce retinal cells following intravitreal AAV delivery in the mouse.^18–20,27–31^ However, relatively few studies have performed systematic analyses of RGC specificity. Among the studies that examined retinal cross sections, one investigated intravitreal delivery of AAV2-CMV on cross sections.^10^ In that study, while Nefh promoter–driven EGFP expression was largely confined to the GCL, CMV-driven EGFP expression extended into the INL, with more than 75% of INL cells expressing EGFP. In addition, many amacrine cells within the GCL also expressed EGFP. Another study directly compared intravitreal injections of AAV2 vectors driven by different promoters, including AAV2-CBA-eGFP, AAV2-CMV-eGFP, AAV2-PGK-eGFP, AAV2-sCAG-eGFP, and AAV2-SYN-eGFP^29^. Consistent with the above findings, the CMV promoter exhibited a largely ubiquitous expression pattern, with transgene expression detected in multiple retinal cell types (45% RBPMS⁺, 25% Prox1⁺, 26% Calretinin⁺, 3% Vimentin⁺, and 1% Calbindin⁺ cells), indicating relatively low RGC specificity. Together, these prior studies are consistent with the results of our current work, demonstrating that the CMV promoter drives transgene expression in a large number of non-RGC retinal cell types following intravitreal AAV delivery. An additional consideration is AAV vector design, which varies across studies and may influence expression patterns. Our AAV vector includes a bovine growth hormone polyadenylation signal (bGH-pA), which is known to enhance transgene expression by improving mRNA stability. It remains unclear whether the AAV vectors used in the studies cited above employed similar or distinct regulatory elements, which could have contributed to differences in overall expression levels and apparent RGC specificity.

This study identified a novel Nrn1-based PEC that enables preferential, age-and injury-resilient gene expression in RGCs via AAV2. Quantification of mGL expression indicated approximately 46% transduction efficiency among RGCs. A more sensitive strategy such as Cre recombinase mediated activation in a Rosa reporter mouse line, where a single Cre event permanently triggers reporter expression, could reveal additional transduced RGC and non-RGCs which could provide a more faithful determination of RGC selectivity.

In present study, we selected three-day post-ONC as an early time point to assess promoter-driven expression largely independent of extensive cell loss. This early time point allows us to evaluate the extent to which injured RGCs express the transgene before massive degeneration occurs. Because studies evaluating neuroprotective gene modulation in RGCs typically administer AAVs prior to or at the time of ONC and analyze retinas at later time points (e.g., 7 or 14 days post-ONC), our experiment provides information on the fraction of RGCs that can be targeted at the onset of injury. Nevertheless, later time points such as 7 days post-ONC, when more than 50% of RGCs are lost are highly relevant for rigorously assessing the utility of this approach. It remains to be determined whether Nrn1 PEC retains its efficacy at these more advanced stages of degeneration.

While this study utilized the wild-type AAV2 capsid, several engineered variants have been reported to enhance retinal gene expression. Mutant AAV2 capsids, including triple or quadruple tyrosine-to-phenylalanine (Y→F) mutants, show improved intracellular trafficking and reduced proteasomal degradation, leading to greater transduction efficiency in retinal neurons.^32,33^ Incorporating the Nrn1 PEC into such optimized capsids could further enhance both the efficiency and consistency of RGC gene expression. Future studies combining the Nrn1 regulatory elements with these AAV capsids may therefore achieve even more robust RGC targeting, broadening their potential for both mechanistic studies and therapeutic applications.

At least three and two Nrn1 isoforms exist in the human and mouse genome, respectively, each driven by distinct regulatory architectures (Supplementary Figs. 2A and 2B). The Nrn1 PEC1 characterized in this study originates from one specific isoform and showed the strongest selectivity for RGC-targeted gene transfer. However, it remains unclear which Nrn1 isoforms are expressed in mouse RGCs, or whether their expression differs across developmental or injury contexts. It is also unknown whether combining PECs from all three isoforms would broaden coverage and enable gene expression to the entire RGC population. Defining isoform usage and testing combinatorial PEC strategies will be important future directions.

While our system demonstrates relatively strong RGC specificity, we see a small fraction of non-RGCs transduced. Although the exact identity of these off-target cells remains to be determined, it is plausible that they include retinal interneurons such as amacrine cells, which reside near RGCs and share some overlapping molecular features.

AAV vectors are constrained by a packaging limit of ∼4.7 kb, which restricts the size of both therapeutic transgenes and their regulatory elements. In this study, while the Nrn1 PEC demonstrated RGC preference and resilience to injury, its minimal functional sequence was not defined. Determining the shortest effective configuration of the Nrn1 PEC may be useful for maximizing available space within the AAV genome. This would allow for the inclusion of larger therapeutic payloads or additional regulatory modules, such as insulators, response elements, or dual-gene systems. Future efforts to streamline the Nrn1 PEC could enhance its versatility as a gene modulation platform for RGCs.

Although our system achieves relatively high preference for RGCs, off-target transduction in non-RGCs may still remain a potential concern, as gene manipulation in even a small fraction of unintended retinal cells could elicit adverse effects. To further refine cell-type specificity and minimize off-target risks, future studies could explore additional Nrn1 PEC variants. This may include constructs with repeated enhancer elements or hybrid configurations incorporating other candidate enhancers. Repeating enhancer sequences has previously been shown to amplify both the strength and precision of gene expression^34^, suggesting that such modifications could yield near-complete RGC specificity while maintaining or improving transduction efficiency.

In summary, the Nrn1 PEC offers a valuable tool for preclinical studies focused on RGCs. Its RGC preference and resilience to injury make it well-suited for investigating RGC biology, neuroprotection, and regeneration in vivo.

## Commercial Relationships Disclosure

None

## SUPPORTING INFORMATION

### Supplementary Figures

**Figure S1:**
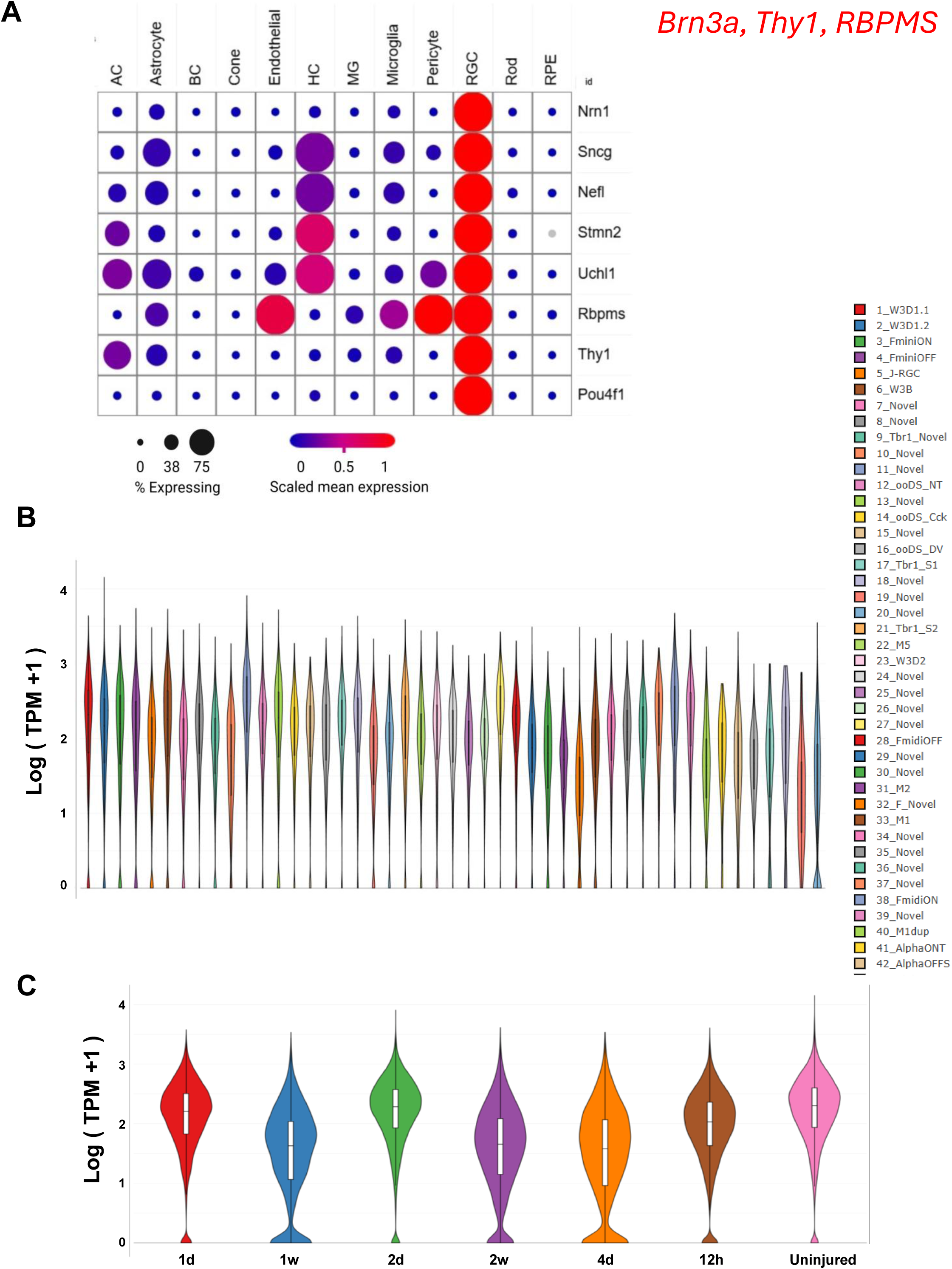
Single cell RNA expression-based profiles of top RGC specific marker genes in adult mouse retina. (**A**) Dot plot showing the scaled mean expression (color intensity) and percentage of cells expressing each gene (dot size) across major retinal cell types. Each column represents a distinct retinal cell population (retinal ganglion cell [RGC], horizontal cell [HC], amacrine cell [AC], microglia, astrocyte, endothelial cell, rod, Müller glia [MG], cone, pericyte, bipolar cell [BC], and retinal pigment epithelium [RPE],). Rows correspond to top five RGC specific genes: *Nrn1, Sncg, Nefl, Stmn2, Uchl1, Rbpms, Thy1* and *Pou4f1.* Red color indicates higher scaled expression, while blue represents lower expression. Larger dot size denotes a higher proportion of cells expressing the gene. Data source.^1^ (**B**) Violin plot showing the expression levels of *Nrn1* across distinct RGC subtypes in healthy mouse retina. Data source.^2^ (**C**) Violin plot depicting *Nrn1* expression in murine RGCs from control (uninjured) retina and retinas at various time points (12 hours, 1, 2, 4-days and 1, 2 – weeks) post-optic nerve crush. Data source.^2^

**Figure S2:**
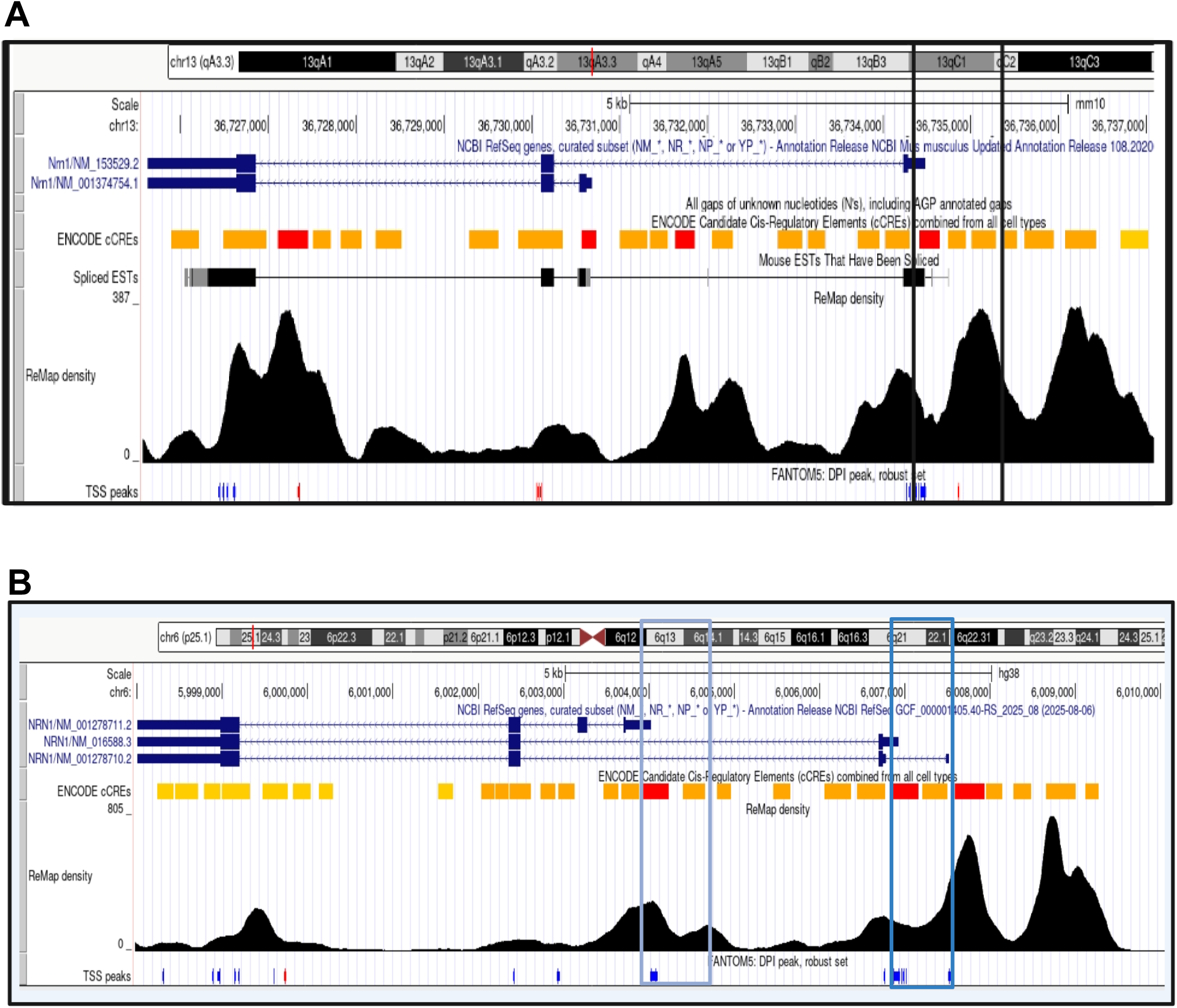
Design and validation of AAV-Nrn1 PEC-driven reporter constructs in adult mouse retina. (**A and B**) UCSC Genome Browser tracks of the Nrn1 genomic regions on mouse (A) and human (B) genome. RefSeq transcripts are shown in blue, with Nrn1 isoforms aligned to annotated exons. Candidate cis-regulatory elements (cCREs) from ENCODE are displayed in red, orange, and yellow, indicating predicted promoter-like signatures, proximal enhancers, and distal enhancer –like signatures, respectively. ReMap density plots highlight transcription factor binding activity, and FANTOM5 CAGE (Cap Analysis of Gene Expression). TSS peaks (blue) mark experimentally validated transcription initiation sites. Vertical rectangle boxes (light blue, blue and black) outline three separate PECs in human and mouse genome selected for synthetic sequence testing. Light blue rectangle represents hNrn1-PEC1; blue rectangle, hNrn1-PEC2; black rectangle, mNrn1-PEC1.

**Figure S3.**
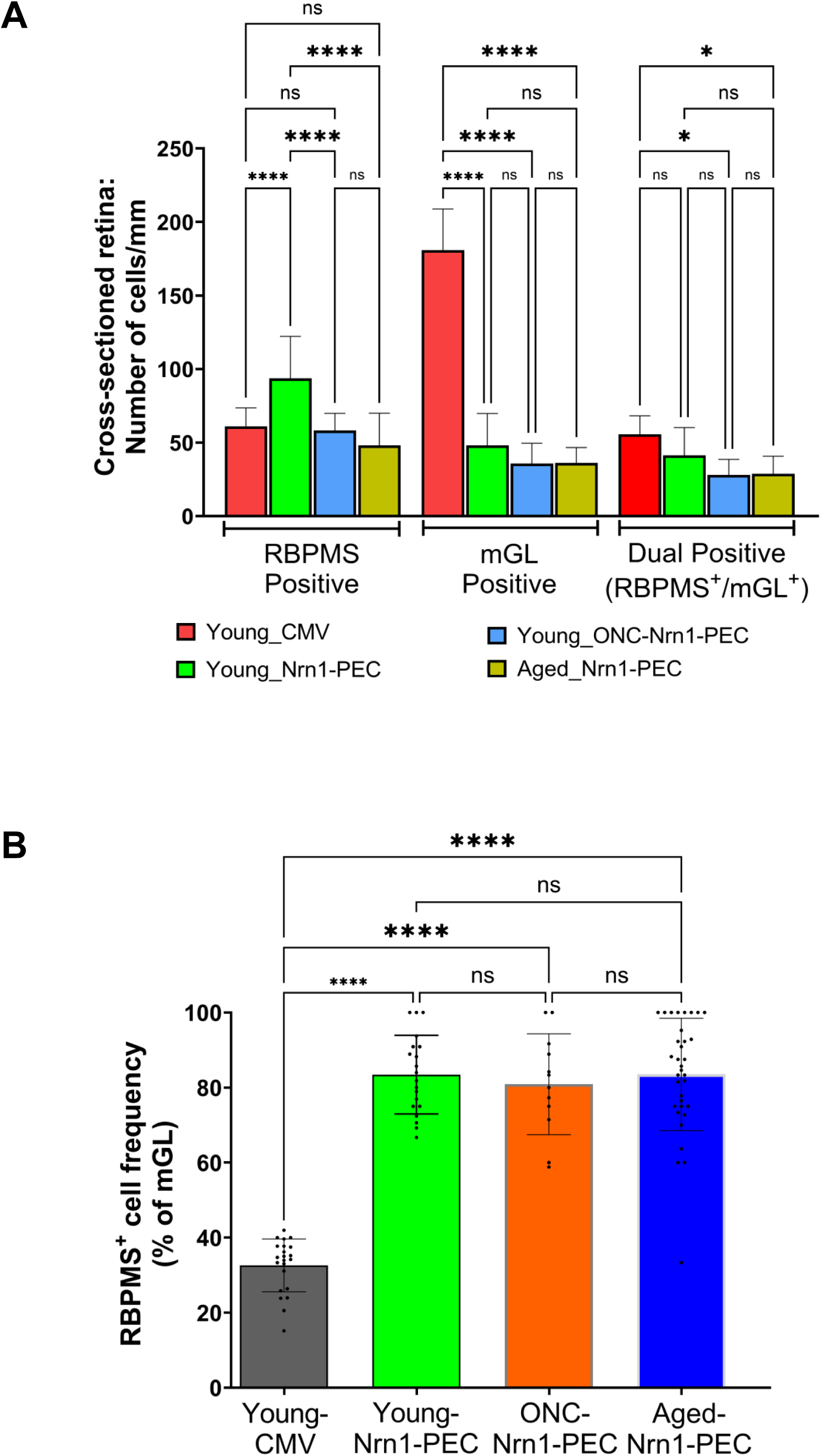
Head-to-head comparison of Nrn1-PEC–driven reporter expression in young, aged, and ONC retina. (**A**) Quantification of RBPMS⁺ cells, mGL⁺ cells, and dual-positive RBPMS⁺/mGL⁺ cells on cross-sectioned mouse retina. Young mice received intravitreal AAV injection driven by either CMV (Young_CMV) or Nrn1-PEC (Young_Nrn1-PEC), while separate cohorts included young mice following optic nerve crush (Young_ONC–Nrn1-PEC) and aged mice receiving Nrn1-PEC (Aged_Nrn1-PEC). (**B**) RBPMS⁺ frequency (% of mGL⁺) as a measure of RGC specificity. Data are presented as mean ± SD (cells/mm). Statistical significance was determined by one-way ANOVA followed by Tukey’s multiple comparisons test; *P < 0.05, ****P < 0.0001; ns, not significant.

